# Synergistic binding of bHLH transcription factors to the promoter of the maize *NADP-ME* gene used in C_4_ photosynthesis is based on an ancient code found in the ancestral C_3_ state

**DOI:** 10.1101/230136

**Authors:** Ana Rita Borba, Tânia S. Serra, Alicja Górska, Paulo Gouveia, André M. Cordeiro, Ivan Reyna-Llorens, Jana Kneřová, Pedro M. Barros, Isabel A. Abreu, M. Margarida Oliveira, Julian M. Hibberd, Nelson J.M. Saibo

## Abstract

C_4_ photosynthesis has evolved repeatedly from the ancestral C_3_ state to generate a carbon concentrating mechanism that increases photosynthetic efficiency. This specialised form of photosynthesis is particularly common in the PACMAD clade of grasses, and is used by many of the world’s most productive crops. The C_4_ cycle is accomplished through cell-type specific accumulation of enzymes but *cis*-elements and transcription factors controlling C_4_ photosynthesis remain largely unknown. Using the *NADP-Malic Enzyme* (*NADP-ME*) gene as a model we aimed to better understand molecular mechanisms associated with the evolution of C_4_ photosynthesis. Two basic Helix-Loop-Helix (bHLH) transcription factors, ZmbHLH128 and ZmbHLH129, were shown to bind the C_4_ *NADP-ME* promoter from maize. These proteins form heterodimers and ZmbHLH129 impairs *trans*-activation by ZmbHLH128. Electrophoretic mobility shift assays indicate that a pair of *cis*-elements separated by a seven base pair spacer synergistically bind either ZmbHLH128 or ZmbHLH129. This pair of *cis*-elements is found in both C_3_ and C_4_ species of the PACMAD clade. Our analysis is consistent with this *cis*-element pair originating from a single motif present in the ancestral C_3_ state. We conclude that C_4_ photosynthesis has co-opted an ancient C_3_ regulatory code built on G-box recognition by bHLH to regulate the *NADP-ME* gene. More broadly, our findings also contribute to the understanding of gene regulatory networks controlling C_4_ photosynthesis.

## Introduction

C_3_ plants inherited a carbon fixation system developed by photosynthetic bacteria, with atmospheric carbon dioxide (CO_2_) being incorporated into ribulose-1,5-bisphosphate (RuBP) by the enzyme Ribulose Bisphosphate Carboxylase/Oxygenase (RuBisCO) to form the three-carbon compound (C_3_) 3-phosphoglycerate (Calvin and Massini 1952). However, RuBisCO can also catalyse oxygenation of RuBP, which leads to the production of 2-phosphoglycolate, a compound that is toxic to the plant cell and needs to be detoxified through an energetically wasteful process called photorespiration (Bowes et al. 1971; Sharkey 1988; Sage 2004). The oxygenase reaction of RuBisCO becomes more common as temperature increases and so in C_3_ plants photorespiration can reduce photosynthetic output by up to 30% (Ehleringer and Monson 1993). In environments such as the tropics where rates of photorespiration are high, C_4_ photosynthesis has evolved repeatedly from the ancestral C_3_ state (Lloyd and Farquhar 1994; Osborne and Beerling 2006). Phylogenetic studies estimate that the first transition from C_3_ to C_4_ occurred around 30 million years ago (MYA) (Christin et al. 2008; Vicentini et al. 2008; Christin et al. 2011). The ability of the C_4_ cycle to concentrate CO_2_ around RuBisCO limits oxygenation and so increases photosynthetic efficiency in conditions where photorespiration is enhanced (Hatch and Slack 1966; Maier et al. 2011; Christin and Osborne 2014; Lundgren and Christin 2016).

The evolution of C_4_ photosynthesis involved multiple modifications to leaf anatomy and biochemistry (Hatch 1987; Sage 2004). In most C_4_ plants, photosynthetic reactions are partitioned between two distinct cell types known as mesophyll (M) and bundle sheath (BS) cells (Langdale 2011). M and BS cells are arranged in concentric circles around veins in the so-called Kranz anatomy (Haberlandt 1904), which enables CO_2_ pumping from M to BS where RuBisCO is specifically located. Atmospheric CO_2_ is first converted to HCO_3_ by carbonic anhydrase (CA) and then combined with phospho*enol*pyruvate (PEP) by PEP-carboxylase (PEPC) to produce oxaloacetate in the M cells. This four-carbon acid (C_4_) is subsequently converted into malate and/or aspartate that transport the fixed CO_2_ from M to BS cells (Kagawa and Hatch 1974; Hatch 1987). Three biochemical C_4_ subtypes are traditionally described based on the predominant type of C_4_ acid decarboxylase responsible for the CO_2_ release around RuBisCO in the BS: NADP-dependent Malic Enzyme (NADP-ME, e.g. *Zea mays*), NAD-dependent Malic Enzyme (NAD-ME, e.g. *Gynandropsis gynandra* formerly designated *Cleome gynandra*) and phospho*enol*pyruvate carboxykinase (PEPCK). However, recent reports suggest that only the NADP-ME and NAD-ME should be considered as distinct C_4_ subtypes, which in response to environmental cues may involve a supplementary PEPCK cycle (Williams et al. 2012; Y. Wang et al. 2014; Rao and Dixon 2016).

The recruitment of multiple genes into C_4_ photosynthesis involved both an increase in their transcript levels (Hibberd and Covshoff 2010) and also patterns of expression being modified from relatively constitutive in C_3_ species (Maurino et al. 1997; Penfield et al. 2004; Taylor et al. 2010; Brown et al. 2011; Maier et al. 2011) to M-or BS-specific in C_4_ plants (Hibberd and Covshoff 2010). Therefore, considerable efforts have been made to identify the transcription factors (TF) and the *cis*-elements they recognise that are responsible for this light-dependent and cell-specific gene expression (Hibberd and Covshoff 2010). Various studies suggest that different transcriptional regulatory mechanisms have been adopted during C_3_ to C_4_ evolution. One is the acquisition of novel *cis*-elements in C_4_ gene promoters that can be recognised by TFs already present in C_3_ plants (Matsuoka et al. 1994; Ku et al. 1999; Nomura et al. 2000), and a second possibility is the acquisition of novel or modified TFs responsible for the recruitment of genes into the C_4_ pathway through *cis*-elements that pre-exist in C_3_ plants (Patel et al. 2006; Brown et al. 2011; Kajala et al. 2012).

A small number of *cis*-elements found in different gene regions have been shown to be sufficient for the M-or BS-specific expression of C_4_ genes. For example, a 41 base pair (bp) Mesophyll Expression Module 1 (MEM1) *cis*-element was identified from the *PEPC* promoter of C_4_ *Flaveria trinervia* and shown to be necessary and sufficient for M cell-specific accumulation of *PEPC* transcripts in C_4_ *Flaveria* species (Gowik et al. 2004). A MEM1-like *cis*-element has also been found in the C_4_ carbonic anhydrase (*CA3*) promoter of *Flaveria bidentis* and shown to drive M cell-specific expression (Gowik et al. 2016). A second *cis-*element named MEM2 and consisting of 9 bp from untranslated regions has also been shown to be capable of directing M-specificity in C_4_ *G. gynandra* (Kajala et al. 2012; Williams et al. 2016). Lastly, in the case of the *NAD-ME* gene from C_4_ *G. gynandra* a region from the coding sequence generates BS-specificity (Brown et al. 2011). In contrast to these insights into *cis*-elements that control cell-specific expression in the C_4_ leaf, no TFs recognising these *cis*-elements have yet been identified.

To address this gap in our understanding, a bottom-up approach was initiated in attempt to identify TFs that regulate the important maize gene *ZmC_4_-NADP-ME* (GRMZM2G085019) that encodes the Malic Enzyme responsible for releasing CO_2_ in the BS cells. Using Yeast One-Hybrid two maize TFs belonging to the superfamily of basic Helix-Loop-Helix (bHLH), ZmbHLH128 and ZmbHLH129, were identified and functionally characterized. In addition, these TFs bind two *cis*-elements synergistically. Analysis of the *cis*-elements in the *NADP-ME* promoters of BEP and PACMAD grass species indicated that this regulation is likely derived from an ancestral G-box that is present in C_3_ species.

## Results

### ZmbHLH128 and ZmbHLH129 homeologs bind FAR1/FHY3 Binding Site *cis*-elements in the *ZmC_4_-NADP-ME* promoter

To identify TFs that interact with the *ZmC_4_-NADP-ME* gene (GRMZM2G085019), we studied the promoter region comprising 1982 base pairs (bp) upstream of the translational start site. This region was divided into six overlapping fragments ranging from 235 to 482 bp in length (supplementary table S1) and used in Yeast One-Hybrid (Y1H). Each fragment was used to generate one yeast bait strain that was then used to screen a maize cDNA expression library. After screening at least 1.3 million colonies for each region of the promoter, two maize bHLH TFs known as ZmbHLH128 and ZmbHLH129 were identified. Both of these TFs bind the promoter between base pairs −389 and −154 in relation to the predicted translational start site of *ZmC_4_-NADP-ME* (fig. 1*A*). These interactions were confirmed by re-transforming yeast bait strains harbouring each of the six sections of the promoter with cDNAs encoding ZmbHLH128 and ZmbHLH129. Consistent with the initial findings, ZmbHLH128 and ZmbHLH129 only activated expression of the *HIS3* reporter when transformed into yeast containing fragment −389 to −154 bp upstream of *ZmC_4_-NADP-ME* (fig. 1*B*, supplementary fig. S1).

**Fig. 1.**
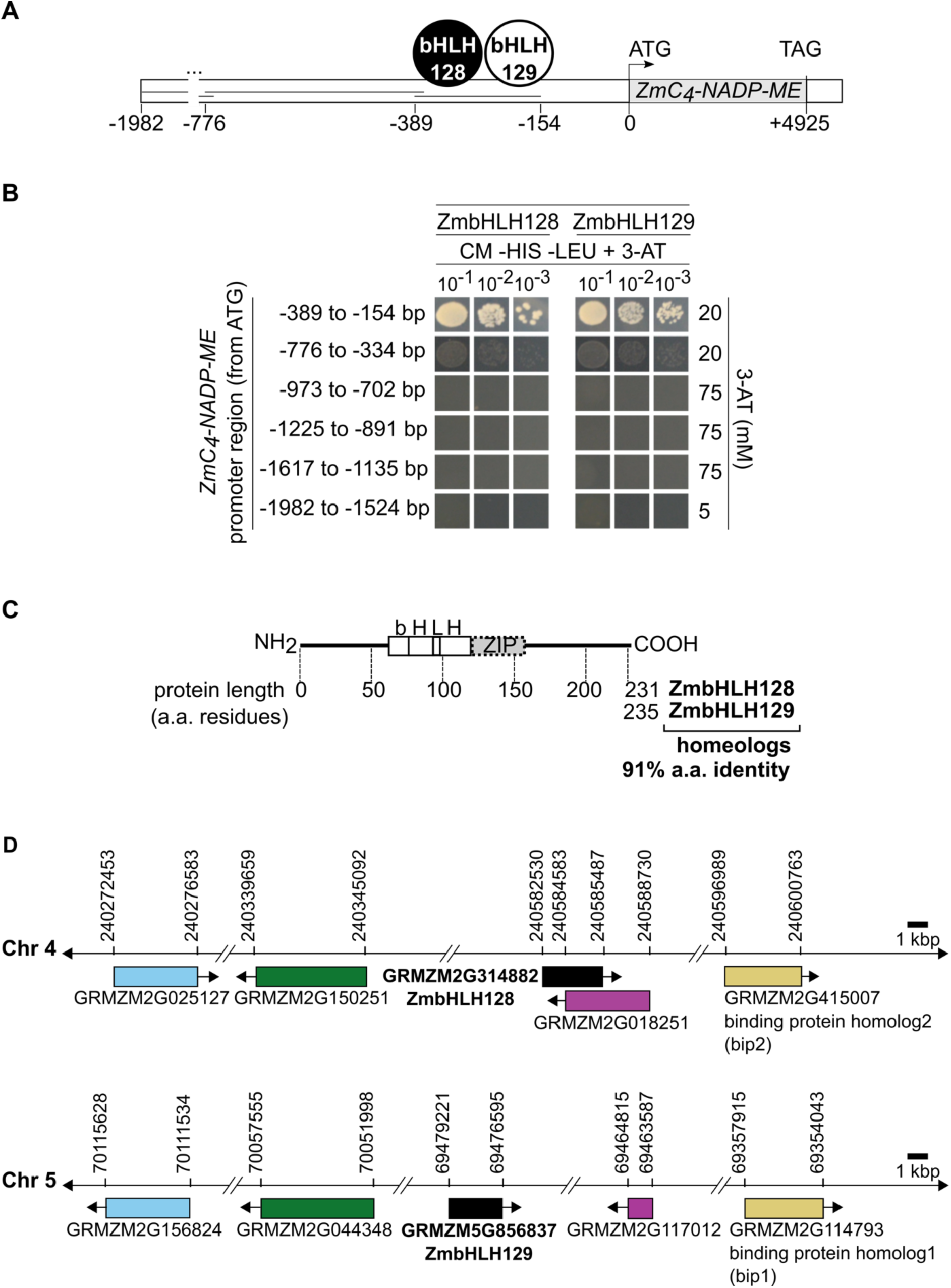
ZmbHLH128 and ZmbHLH129 homeologs bind the *ZmC_4_-NADP-ME* promoter. (*A*) Schematic representation of the *ZmC_4_-NADP-ME* promoter, divided into fragments used as baits in Y1H screenings, and the ZmbHLH TFs identified. ATG and TAG are the translational start codon and the stop codon of the *ZmC_4_-NADP-ME* ORF, respectively. ZmbHLH position on the scheme indicates that they bind between the base pairs −389 and −154 in relation to the ATG. (*B*) Analysis of ZmbHLH-*pZmC_4_-NADP-ME* binding specificity. Each of the six yeast bait strains was transformed with both ZmbHLHs (pAD-GAL4-2.1∷TF vectors) and positive interactions selected on CM-HIS-LEU + 3-AT (yeast Complete Minimal medium lacking histidine and leucine amino acids, and supplemented with 3-amino-1,2,4-triazole (3-AT), a competitive inhibitor of the *HIS3* gene product). (*C*) Schematic representation of basic Helix-Loop-Helix (bHLH) and leucine zipper (ZIP) protein domains, and respective position in protein sequences. (*D*) Schematic representation of *ZmbHLH128* and *ZmbHLH129* (black) and four additional maize homeolog gene pairs located in syntenic regions of chromosomes 4 and 5. Homeolog genes are indicated by colour. Arrows indicate direction of transcription of each gene. Genomic coordinates provided from the B73 RefGen_v3 assembly version.

ZmbHLH128 and ZmbHLH129 possess a bHLH domain followed by a contiguous leucine zipper (ZIP) motif (fig. 1*C*). This bHLH domain is highly conserved between both ZmbHLHs and consists of 61 amino acids that can be separated into two functionally distinct regions. The first is a basic region located at the N-terminal end of the bHLH domain and is involved in DNA binding, and the second is a Helix-Loop-Helix region mediating dimerization towards the carboxy-terminus (fig. 1*C*) (Murre et al. 1989; Toledo-Ortiz et al. 2003). ZmbHLH128 and ZmbHLH129 share 91% amino acid identity (fig. 1*C*) and they are encoded by homeolog genes located in syntenic regions of maize chromosomes 4 and 5 (fig. 1*D*, supplementary table S2).

Although ZmbHLH128 and ZmbHLH129 both possess three amino acids involved in G-box binding (K9, E13, and R17) (Massari and Murre 2000; Li et al. 2006), this family of TFs has also been shown to bind to N-box (5’-CACGCG-3’), N-box B (5’-CACNAG-3’) and FBS (FAR1/FHY3 Binding Site, 5’-CACGCGC-3’) motifs (Sasai et al. 1992; Ohsako et al. 1994; Fisher and Caudy 1998; Kim et al. 2016). Therefore, the *ZmC_4_-NADP-ME* promoter was assessed for additional *cis*-elements to which ZmbHLH128 and ZmbHLH129 might bind. A total of eight such *cis*-elements were found, consisting of two N-boxes B, two N-boxes, one G-box, two FBSs and one E-box (fig. 2*A*). Electrophoretic Mobility Shift Assays (EMSA) were used to test whether ZmbHLH128 and ZmbHLH129 were able to interact with each of these *cis-*elements *in vitro* (fig. 2*B* and *C*). Consistent with the Y1H findings, EMSA showed that recombinant Trx∷ZmbHLH128 and Trx∷ZmbHLH129 proteins caused an uplift of radiolabeled probes containing FBS *cis*-elements (probes 6, 7, and 6+7) (fig. 2*C*), positioned between nucleotides −389 and −154 in relation to the predicted translational start site (see fig. 1*A*). ZmbHLH128 also showed weak binding to probe 3 that contained a N-box *cis*-element that was not bound by ZmbHLH128 or ZmbHLH129 in Y1H (see fig. 1*B*), and signal intensity was similar to that observed from probe 7 (fig. 2*C*). It is possible that relatively weak binding to probe 7 is due to it being three nucleotides-shorter than the other probes (fig. 2*B*). Trx alone and OsPIF14 (a bHLH known to bind the N-box motif (Cordeiro et al. 2016)) were used as negative controls (fig. 2*C*). The two FBS motifs, in probe 6+7, are separated by a short 7 bp spacer sequence and are found in opposite orientations (fig. 2*B*). The increase in band intensities detected when both *cis-*elements were combined (fig. 2*C*) suggests that they may function synergistically. Overall, these data indicate that ZmbHLH128 and ZmbHLH129 target 21bp of DNA sequence (7bp FBS, 7bp spacer, and 7bp FBS).

**Fig. 2.**
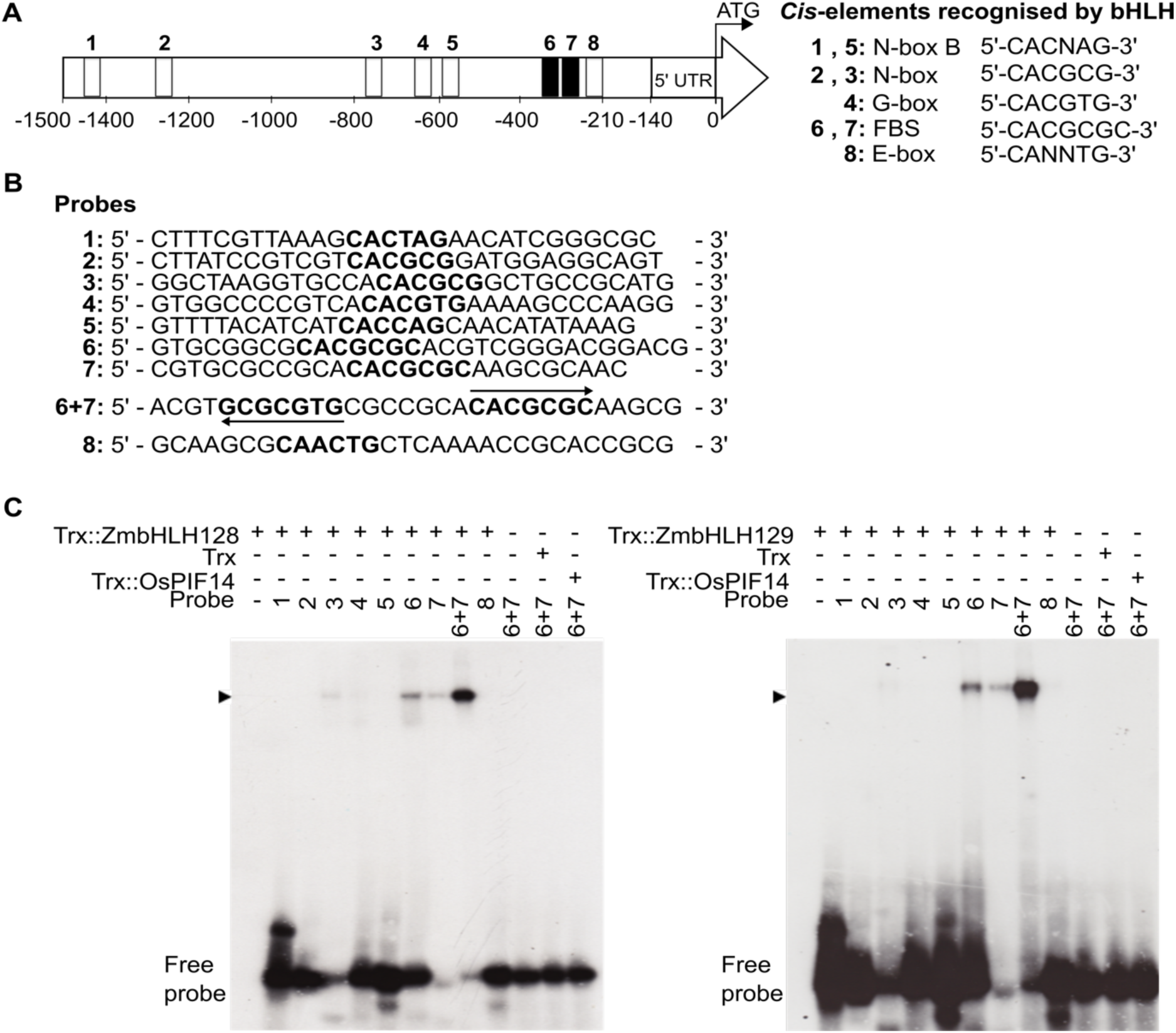
ZmbHLH128 and ZmbHLH129 bind two FBS *cis*-elements present in *ZmC_4_-NADP-ME* promoter. (*A*) Schematic representation of position and nucleotide sequence of eight *cis*-elements recognised by bHLH that were identified in the *ZmC_4_-NADP-ME* promoter. FBS stands for FHY3/FAR1 Binding Site and it is a N-box-containing motif. (*B*) EMSA probe sequences used to test *in vitro* binding affinity of ZmbHLH128 and ZmbHLH129 to *cis*-elements (highlighted in bold). Arrows indicate that the FBS *cis*-elements are present in opposite orientations. (*C*) EMSAs showing *in vitro* binding affinity of Trx∷ZmbHLH128 (gel on the left) and Trx∷ZmbHLH129 (gel on the right) to the radiolabeled probes described in (*B*). Arrowheads indicate uplifted ZmbHLH-DNA probe complexes. Free probe indicates unbound DNA probes.

### ZmbHLH128 and ZmbHLH129 form both homo-and heterodimers and ZmbHLH129 impairs *trans*-activation by ZmbHLH128

Because ZmbHLH128 and ZmbHLH129 bind the FBS *cis*-elements in close proximity but also possess domains mediating protein dimerization, we next investigated whether these proteins form homo-and/or heterodimers. *In vitro*, the recombinant Trx∷ZmbHLH128 and Trx∷ZmbHLH129 proteins formed homodimers (fig. 3*A*). To confirm this interaction *in vivo*, as well as to test for heterodimerization, Bimolecular Fluorescence Complementation Assays (BiFC) in maize protoplasts were performed. Whilst negative controls produced no YFP fluorescence, ZmbHLH128 and ZmbHLH129 formed both homo-and heterodimers (fig. 3*B*). With the exception of ZmbHLH129 homodimers whose location extended to the cytoplasm and plasma membrane, in each case YFP signal was specifically localised to the nucleus (fig. 3*B*). Nuclear localisation of these ZmbHLH proteins supports their roles as transcriptional regulators.

**Fig. 3.**
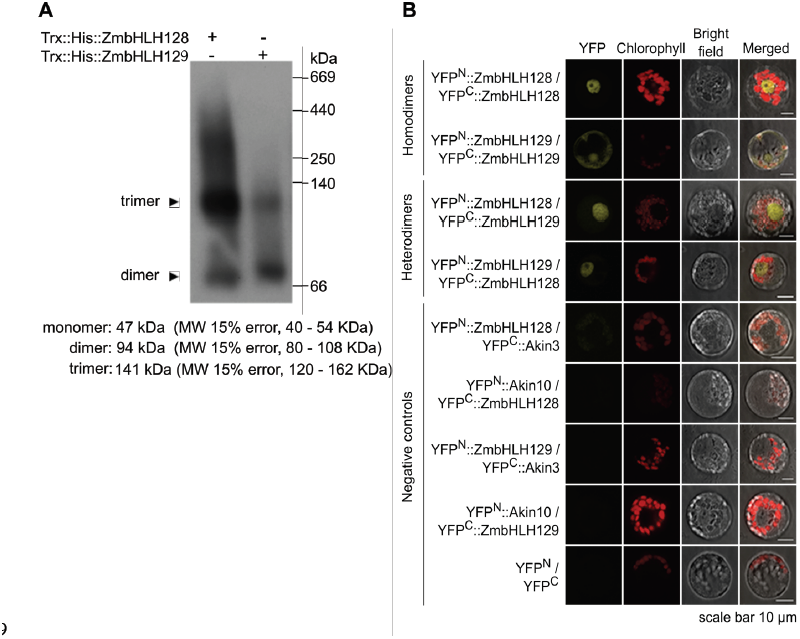
ZmbHLH128 and ZmbHLH129 form both homo-and heterodimers. (*A*) Western blot of BN-PAGE for the recombinant proteins Trx∷His∷ZmbHLH128 and Trx∷His∷ZmbHLH129. Gel was loaded with equivalent amount of protein. Recombinant proteins were immunodetected using α-His antibody. MW indicates molecular-weight size marker. (*B*) Protein interactions between ZmbHLH128 and ZmbHLH129 were tested by BiFC in maize mesophyll protoplasts co-transformed with constructs expressing ZmbHLH128 and ZmbHLH129 fused to N-and C-terminal YFP domains. YFP^N^ and YFP^C^ indicate split N-and C-terminal YFP domains, respectively.

To test the capacity of ZmbHLH128 and ZmbHLH129 to regulate transcription, transient expression assays were performed in leaves of *Nicotiana benthamiana*. The *GUS* reporter gene driven by the fragment of *pZmC_4_-NADP-ME* to which ZmbHLH128 and ZmbHLH129 bind was used as reporter, whilst the full-length *ZmbHLH128* and *ZmbHLH129* CDS sequences driven by the constitutive *CaMV35S* promoter were used as effectors (fig. 4*A*). Co-infiltration of this reporter with the ZmbHLH128 effector resulted in an increase in GUS activity, indicating that ZmbHLH128 can act as a transcriptional activator (fig. 4*B*). In contrast, ZmbHLH129 showed no intrinsic *trans*-activation activity (fig. 4*C*). In order to test whether the ZmbHLH128-ZmbHLH129 heterodimers had a different *trans*-activation activity from ZmbHLH128 or ZmbHLH129 homodimers, leaves were co-infiltrated with the reporter and both effectors simultaneously. Interestingly, the *trans*-activation activity observed for the ZmbHLH128 alone (fig. 4*B*) was lost when this TF was co-expressed with its homeolog ZmbHLH129 (fig. 4*D*).

**Fig. 4.**
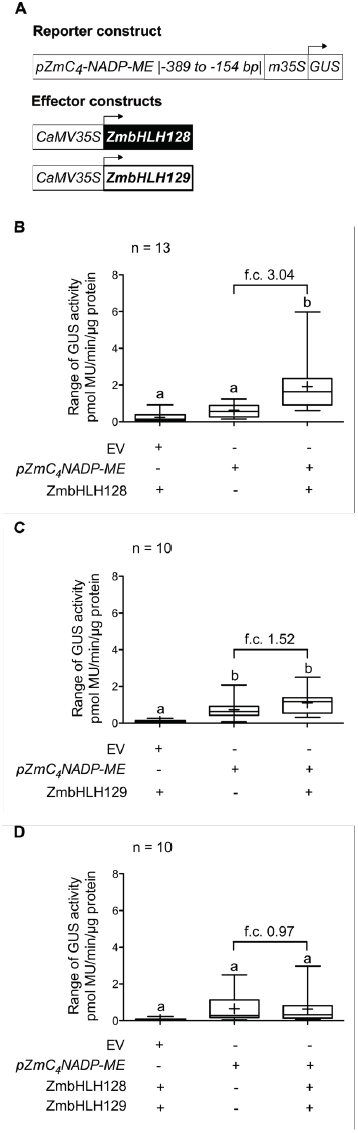
ZmbHLH129 impairs *trans*-activation of the *ZmC_4_-NADP-ME* promoter by ZmbHLH128. (*A*) Schematic representation of reporter and effector constructs used in transient expression assays in leaves of *Nicotiana benthamiana*. Reporter construct contains *GUS* gene driven by the minimal *CaMV35S* promoter (m35S) fused to *pZmC_4_-NADP-ME* (-389 to −154 bp). Effector constructs contain the *ZmbHLH128* or *ZmbHLH129* CDS driven by the full *CaMV35S* promoter. (*B*-*D*) Box plots (2.5 to 97.5 percentiles) showing GUS activity, expressed in picomoles of the reaction product 4-methylumbelliferone (MU) generated per minute per microgram of protein, in leaves agro-infiltrated with reporter and the following effector constructs: (*B*) ZmbHLH128, (*C*) ZmbHLH129, and (*D*) ZmbHLH128 and ZmbHLH129. Different letters denote differences in experimental data that are statistically significant (One-way ANOVA, Tukey test, *p* ≤ 0.05, n = 10-13). EV indicates pGWB3i empty vector (no promoter fragment cloned). Cross inside box plots indicates mean. f.c. indicates fold-change.

### The G-box-based *cis*-element pair recognised by ZmbHLH128 and ZmbHLH129 in *NADP-ME* promoters operates synergistically

To understand whether the two FBS *cis*-elements identified in the promoter of *ZmC_4_-NADP-ME* (see fig. 2) are associated with the evolution of C_4_ photosynthesis, we investigated whether they are conserved in promoters of other *NADP-MEs* from C_3_ and C_4_ grass species. Three C_3_ species (*Dichanthelium oligosanthes*, *Oryza sativa* and *Brachypodium distachyon*) and three C_4_ species (*Zea mays*, *Sorghum bicolor* and *Setaria italica*) were assessed (fig. 5*A*). Within the C_4_ species, *Zea mays* and *Sorghum bicolor* possess two plastidic NADP-ME isoforms: one that is used in C_4_ photosynthesis (C_4_-NADP-ME, GRMZM2G085019 and Sobic.003g036200) and a second one not involved in the C_4_ cycle (*non*C_4_-NADP-ME, GRMZM2G122479 and Sobic.009g108700). In contrast, *S. italica* possesses only one plastidic NADP-ME isoform that is used in the C_4_ cycle (C_4_-NADP-ME, Si000645) (Alvarez et al. 2013).

**Fig. 5.**
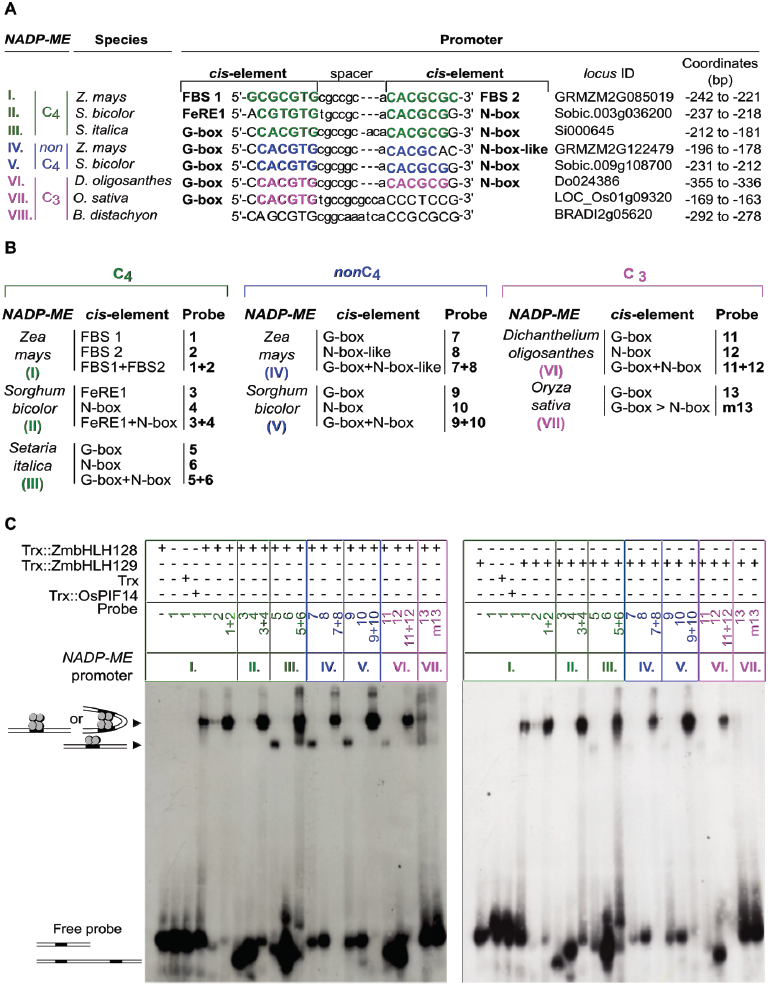
The G-box-based *cis-*element pair recognised by ZmbHLH128 and ZmbHLH129 in *NADP-ME* promoters operates synergistically. (*A*) Sequence alignment of the two FBS *cis*-elements present in *ZmC_4_-NADP-ME* promoter against homologous *cis*-elements present in other promoters of genes encoding plastidic NADP-ME. C_4_ grasses: *Zea mays*, *Sorghum bicolor* and *Setaria italica*; C_3_ grasses: *Dichanthelium oligosanthes*, *Oryza sativa* and *Brachypodium distachyon*. Plastidic NADP-MEs are colour-coded: green for C_4_, blue for *non*C_4_ and magenta for C_3_. *Cis*-elements are highlighted in bold and coloured according to the NADP-ME they belong to. FBS stands for FHY3/FAR1 Binding Site and FeRE1 for Iron Responsive Element 1. (*B*) EMSA probes used to test *in vitro* binding affinity of ZmbHLH128 and ZmbHLH129 to each *cis*-element described in (*A*). Probe sequences are listed in supplementary table S3. (*C*) EMSA assays showing *in vitro* binding affinity of Trx∷ZmbHLH128 (gel on the left) and Trx∷ZmbHLH129 (gel on the right) proteins to the probes described in (*B*). Arrowheads indicate uplifted ZmbHLH-DNA probe complexes. Free probe indicates unbound DNA probes.

Although in C_3_ *B. distachyon* no homologous *cis*-elements to the FBSs in the *ZmC_4_-NADP-ME* promoter were detected, in *O. sativa* one G-box was found in the same position as FBS 1 from *Z. mays*. Moreover, in the other promoters, *cis*-elements that can bind bHLH proteins were present in pairs (fig. 5*A*). In both the C_3_ and C_4_ grasses these *cis*-element pairs flank a spacer that is highly conserved in sequence and length (7 to 9 bp) (fig. 5*A*). The C_4_-*NADP-ME* promoters from *Z. mays* and *S. bicolor* share a common mutation in the third nucleotide position of the alignment (A?G) (fig. 5*A*). Two additional mutations are specific to *Z. mays* (the first and last nucleotides of FBS 1 and FBS 2, respectively), whilst one is *S. bicolor*-specific (C?T at the fourth position) (fig. 5*A*). It is possible that mutations unique to *Z. mays* or *S. bicolor* are neutral and the main impact on C_4_-*NADP-ME* gene expression is due to mutation in the third nucleotide in the common ancestor of *Z. mays* and *S. bicolor*. Alternatively, it is also possible that both this mutation in the last common ancestor and species-specific modifications impacted on gene expression of C_4_-*NADP-ME*.

To test if ZmbHLH128 and ZmbHLH129 bind the *cis*-elements identified from these additional species EMSA was performed on each *cis*-element separately as well as the *cis*-element pairs found in each *NADP-ME* promoter (fig. 5*B* and *C*, supplementary table S3). ZmbHLH128 and ZmbHLH129 showed low binding affinity for the single G-box identified in the *O. sativa* promoter (probe 13) and binding affinity was not increased by mutating the G-box to a canonical N-box (probe m13) (fig. 5*B* and *C*). This low binding affinity behaviour for single G-box *cis*-elements was consistent for all the *NADP-ME* promoters containing G-boxes (probes 5, 7, 9 and 11) (fig. 5*B* and *C*). Although both ZmbHLHs did not show binding affinity for the additional N-boxes or N-box-like alone (probes 6, 8, 10 and 12) (fig. 5*B* and *C*), when these additional motifs were acquired and formed a pair with the ancestral G-box, binding affinity was increased (probes 5+6, 7+8, 9+10 and 11+12) and led to an increased uplift compared with the G-boxes alone (probes 5, 7, 9 and 11) (fig. 5*B* and *C*). Given the similar length of probes 1, 2, 1+2, 5, 7, 9 and 11 (24 to 30 bp) (supplementary table S3), it is possible that this difference in migration of ZmbHLH-probe complexes results from the binding of bHLH to G-boxes in a lower oligomeric state (supplementary fig. S2), which based on the literature must be dimers (De Masi et al. 2011). Strong binding of *cis*-element pairs was also observed when the ancestral G-box evolved into either FBS or FeRE1 elements found in C_4_ *Z. mays* and *S. bicolor* (probes 1+2 and 3+4) (fig. 5*B* and *C*). In the C_4_ *Z. mays* promoter, both ZmbHLHs showed binding affinity for single FBS *cis*-elements (probes 1 and 2) in the highest oligomeric state (fig. 5*B* and *C*, supplementary fig. S2).

Since ZmbHLH128 and ZmbHLH129 showed weak binding to single *cis*-elements, we tested their binding by mutating these *cis*-elements in probes with the pairs (supplementary fig. S3). For each pair, three mutant probes were designed: two in which the two *cis*-elements were mutated individually (keeping one *cis*-element wild-type) and one in which both *cis*-elements were mutated simultaneously (supplementary table S3). Competition experiments were performed using radiolabeled wild-type probes (with *cis*-element pairs) and 200- to 400-fold excess of unlabeled wild-type and mutant probes (supplementary fig. S3). Binding of both ZmbHLHs to the labeled wild-type probes could be efficiently out-competed by unlabeled wild-type and mutant probes in which the following *cis*-elements were not mutated: FBS 1 (in *Z. mays* C_4_-*NADP-ME*, probe 1+m2-A, supplementary fig. S3*A*); FBS 2 (in *Z. mays* C_4_-*NADP-ME*, probe m1+2-B, supplementary fig. S3*A*); N-box (in *S. bicolor* C_4_-*NADP-ME*, probe m3+4-E, supplementary fig. S3*B*); and G-box (in *S. italica* C_4_-*NADP-ME*, probe 5+m6-G, supplementary fig. S3*C*; *Z. mays non*C_4_-*NADP-ME*, probe 7+m8-J, supplementary fig. S3*D*; *S. bicolor non*C_4_-*NADP-ME*, probe 9+m10-M, supplementary fig. S3*E*; and *D. oligosanthes* C_3_-*NADP-ME*, probe 11+m12-P, supplementary fig. S3*F*). These EMSA competition experiments thus confirmed binding of ZmbHLH128 and ZmbHLH129 to the *cis*-elements described above. Taken together, the results indicate that a second *cis*-element recognised by bHLH TFs is acquired in the promoters of genes encoding plastidic NADP-ME and that each *cis*-element pair operates synergistically to allow interaction with either ZmbHLH128 or ZmbHLH129 in C_3_ and C_4_ grasses (fig. 5, supplementary fig. S2 and S3).

Given the binding affinity *in vitro* of ZmbHLH128 and ZmbHLH129 to the G-box in the *ZmnonC_4_-NADP-ME* promoter (probes 7 and 7+8, fig. 5*C*), we tested their binding ability *in planta*. Transient expression assays were performed in leaves of *N. benthamiana* co-infiltrated with *GUS* reporter gene driven by a *ZmnonC_4_-NADP-ME* promoter fragment containing the *cis*-element pair G-and N-box-like (-368 to −143 bp) and the effector constructs ZmbHLH128 and ZmbHLH129 (supplementary fig. S4*A*). Compared with the reporter alone, co-infiltration of *ZmnonC_4_-NADP-ME* reporter and the ZmbHLH128 and ZmbHLH129 effectors did not impact on GUS activity in tobacco system (supplementary fig. S4*B*-*D*). These results suggest that although ZmbHLH128 on its own binds both the *ZmC_4_-NADP-ME* and *ZmnonC_4_-NADP-ME* promoters *in vitro* (probes 1, 2, 1+2, 7 and 7+8, fig. 5*B* and *C*), this might not be the case *in planta* (supplementary fig. S4).

### Acquisition of N-box-derived *cis*-elements in *NADP-ME* promoters facilitates ZmbHLH128 and ZmbHLH129 binding in PACMAD grasses

Phylogenetic analysis of the genes encoding C_3_ and C_4_ plastidic NADP-MEs reflects previously reported grass species phylogeny (fig.6*A*) (Grass Phylogeny Working Group II 2012). It inferred two main clades: one formed by C_3_ BEP species (*B. distachyon* and *O. sativa*) and a second formed by C_3_ (*D. oligosanthes*) and C_4_ PACMAD species (*S. italica*, *S. bicolor* and *Z. mays*) (fig.6*A*).

**Fig. 6.**
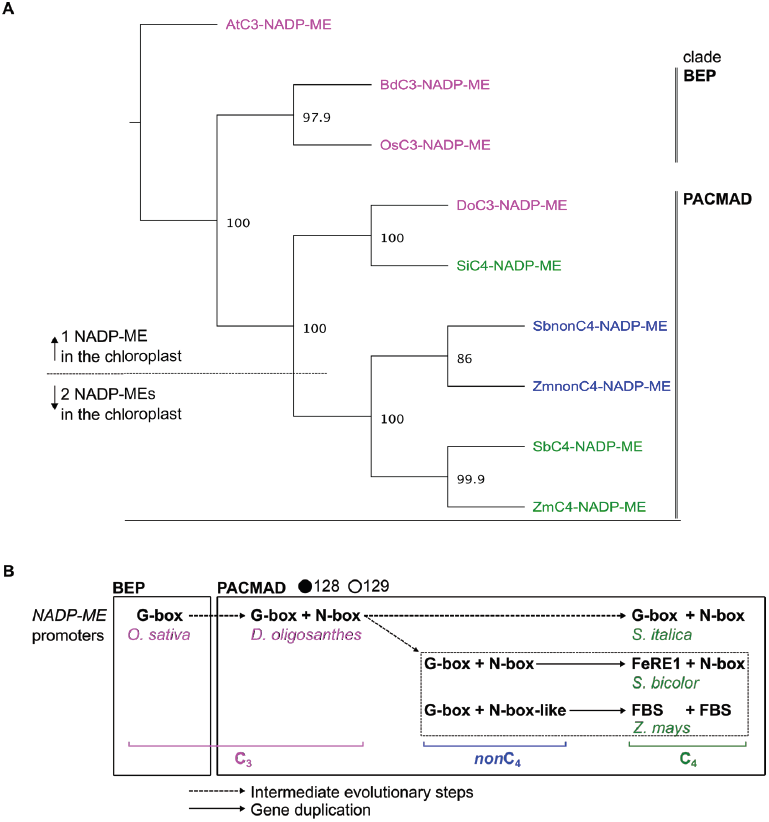
Acquisition of N-box-derived *cis*-elements in *NADP-ME* promoters facilitates ZmbHLH128 and ZmbHLH129 binding in PACMAD grasses. (*A*) Phylogenetic tree of genes encoding plastidic NADP-ME from C_3_ and C_4_ grass species. C_3_: *Brachypodium distachyon (Bd), Oryza sativa* (Os) and *Dichanthelium oligosanthes* (Do); C_4_: *Setaria italica* (Si), *Sorghum bicolor* (Sb) and *Zea mays* (Zm). *NADP-ME*s are colour-coded: magenta for C_3_, blue for *non*C_4_ and green for C_4_. *NADP-ME* genomic sequences were aligned using MUSCLE, and the phylogenetic tree inferred by NJ method (1000 bootstrap pseudoreplicates, node numbers indicate bootstrap values). Gene encoding C_3_ plastidic NADP-ME from *Arabidopsis thaliana* (AtC_3_-NADP-ME) was used as outgroup. (*B*) Diagram representing C_3_ to C_4_ molecular evolution of homologous bHLH binding *cis*-elements identified in promoters of genes encoding plastidic NADP-ME. Black and white circles represent ZmbHLH128 and ZmbHLH129 binding ability to *NADP-ME* gene promoters, respectively.

Based on the observed nucleotide modifications in *cis-*elements recognised by bHLH TFs, we propose a model relating to the recruitment of *NADP-ME* into C_4_ photosynthesis in grasses (fig. 6*B*). This proposes that an ancestral G-box found in the *NADP-ME* promoter from C_3_ BEP *O. sativa* was conserved throughout C_3_ to C_4_ evolution and is shared by different C_3_ and C_4_ grass lineages. However, in the PACMAD group a second *cis*-element recognised by bHLH was acquired such that the *NADP-ME* gene from the C_3_ species *D. oligosanthes* and genes encoding *non*C_4_-NADP-ME from C_4_ *S. bicolor* and *Z. mays* all contain a G-and N-box/N-box-like pair. In C_4_ *S. italica* this *cis*-code has been retained in the C_4_-*NADP-ME*, but in *S. bicolor* and *Z. mays* the original G-box has evolved to become either a FeRE1 or a FBS element, respectively (fig. 6*B*). Overall, these results suggest that the acquisition of N-box-derived *cis*-elements may have facilitated ZmbHLH128 and ZmbHLH129 binding to promoters of genes encoding plastidic NADP-ME in the PACMAD clade.

## Discussion

### ZmbHLH128 and ZmbHLH129 homeologs interact with maize C_4_-and *non*C_4_-*NADP-ME* promoters *in vitro* showing different *trans*-activation activity *in planta*

In this study, we showed that ZmbHLH128 and ZmbHLH129 form a maize homeolog pair resulting from the recent maize whole genome duplication (WGD) event that occurred 5-12 million years ago. This WGD occurred ~25 million years after C_4_ photosynthesis evolved in the Chloridoideae subfamily of the PACMAD clade (Christin et al. 2008; Christin et al. 2011). As the length of exons 1 and 2 and the total number of amino acids in the mature protein of ZmbHLH128 are more similar to sorghum ortholog SbbHLH66 (supplementary fig. S5), we propose that ZmbHLH129 has diverged more from the ancestral gene. Both of these TFs bind two FBS *cis*-elements that are in close proximity in the maize C_4_*-NADP-ME* (GRMZM2G085019) promoter. Although ZmbHLH128 has been predicted *in silico* to regulate C_4_ photosynthesis (L. Wang et al. 2014), as far as we are aware, this is the first report of its functional characterization. ZmbHLH128 alone activates *ZmC_4_-NADP-ME* gene expression, whilst ZmbHLH129 alone shows no *trans*-activation activity on this promoter. As the duplication event that generated ZmbHLH129 took place after the evolution of C_4_ photosynthesis, it seems possible that this gene is not required for C_4_ photosynthesis. ZmbHLH128 and ZmbHLH129 form heterodimers and despite ZmbHLH128 activating the expression of *ZmC_4_-NADP-ME* its regulatory activity is impaired by its homeolog ZmbHLH129. To explain this impairment, we hypothesise different scenarios that may occur *in vivo*: either ZmbHLH128 and ZmbHLH129 act as heterodimers and ZmbHLH128 loses its DNA binding activity when combined with ZmbHLH129 or they act as homodimers and compete directly for the same FBSs, towards which ZmbHLH129 has a higher binding affinity. The former scenario has been described for bZIP TFs from *Arabidopsis*, where bZIP63 has negative effects on the formation of bZIP1-DNA complexes probably due to conformational differences between bZIP1 homodimer and bZIP1-bZIP63 heterodimers (Kang et al. 2010). The latter scenario has been reported for the maize Dof1 and Dof2 TFs. Dof1 is a transcriptional activator of light-regulated genes in leaves, however, in stems and roots, this TF is not able to regulate those genes since the repressor Dof2 is expressed there and blocks Dof-specific *cis*-elements (Yanagisawa and Sheen 1998).

In addition to the capacity of ZmbHLH128 and ZmbHLH129 to interact with FBSs found in the maize C_4_-*NADP-ME* promoter, both ZmbHLHs were shown to bind *in vitro* to the promoter of maize *non*C_4_*-NADP-ME* (GRMZM2G122479) that possesses the *cis*-element pair G-and N-box-like. *In planta*, however, ZmbHLH128 and ZmbHLH129 showed no *trans*-activation activity on this promoter. It is well known that primary DNA sequence and its structural properties are determinants of DNA binding specificity *in vivo* (Rohs et al. 2009) and so it is possible that both ZmbHLHs display increased *in vivo* binding specificity for the FBS pair in the *ZmC_4_-NADP-ME* promoter than for the G-and N-box-like pair in the *ZmnonC_4_-NADP-ME* promoter. Therefore, ZmbHLH128 seems to affect the level of expression of *NADP-ME* as it activates the *ZmC_4_-NADP-ME* promoter through the pair formed by two FBSs but the same trend was not observed for the *ZmnonC_4_-NADP-ME* promoter with the G-and N-box pair. Additionally, we hypothesise that these modifications of promoter sequences may also affect light/circadian regulation of the *ZmC_4_-NADP-ME* gene as FBS *cis*-elements have been described in promoters of circadian-clock-regulated and light-responsive genes (Lin et al. 2007; Li et al. 2011; Kim et al. 2016). The mutation of two close FBSs in the promoter of the circadian-clock gene EARLY FLOWERING 4 (*ELF4*) proved to be sufficient to abolish its rhythmic expression (Li et al. 2011). More broadly, our findings also contribute to the understanding of gene regulatory networks controlling C_4_ photosynthesis.

### The G-box-based *cis*-element pair present in *NADP-ME* promoters synergistically bind either ZmbHLH128 or ZmbHLH129

We identified a *cis*-element pair recognised by bHLH that occupy homologous positions in *NADP-ME* promoters from C_3_ and C_4_ grasses. These *cis*-elements flank a short spacer and operate synergistically to facilitate interaction with ZmbHLH128 and ZmbHLH129. We suggest a mechanism by which these TFs may be recruited to the *cis*-elements associated with C_4_ photosynthesis. We propose that one *cis*-element is sufficient to recruit a bHLH homodimer (G-box) or tetramer (N-box or FBS in promoters where the ancestral G-box is no longer present), however, the presence of a second *cis*-element in the vicinity increases bHLH binding affinity (supplementary fig. S2). It is possible that both *cis*-elements are brought together through the interaction with a bHLH tetramer formed by two dimers, which may involve DNA bending (supplementary fig. S2). Therefore, this *cis*-element pair could operate synergistically to confer stabilisation of bHLH binding. This mechanism of TF-DNA assembly has previously been proposed for MADS-domain TFs that can bind two nearby CArG boxes through DNA looping and formation of tetrameric complexes (Theissen 2001; Theissen and Saedler 2001; Melzer and Verelst 2009; Smaczniak et al. 2012; Smaczniak et al. 2017). In this case, and consistent with our results, MADS-domain TFs were found to bind single CArG boxes either as dimers or tetramers, however, when their target gene promoters contain CArG box pairs they bind as tetramers (Smaczniak et al. 2012). It has been proposed that the probability of DNA loop formation increases with shorter distances between *cis*-elements due to the low elastic bending energy required to bring the protein dimers together (Agrawal et al. 2008). Interestingly, in all *NADP-ME* promoters assessed in this study except rice and Brachypodium the two *cis*-elements were found to be in close proximity, which may encourage DNA looping. In addition to the spacer length, its sequence appears highly conserved. This is consistent with evidence suggesting that nucleotides outside core *cis*-elements affect TF binding specificity by providing genomic context and influencing three-dimensional structure (Atchley et al. 1999; Martínez-Garcia et al. 2000; Grove et al. 2009; Gordân et al. 2013). For example, Cbf1 and Tye7 are yeast bHLHs that show preference for a subset of G-boxes present throughout the yeast genome (Gordân et al. 2013). These differences in binding preferences were observed not just *in vivo* but also *in vitro* and so DNA sequences flanking core G-boxes were found to explain this differential bHLH-G-box binding (Gordân et al. 2013).

The mechanism proposed here for how bHLH TFs interact with their target *cis*-elements suggests that these DNA sequences are not randomly arranged in gene promoters and may affect how *cis*-element specificity is achieved. Indeed, in some promoters bound by bHLH TFs two or more *cis*-elements were found to be clustered. For example, two overlapping FBSs were reported in the 400 base pairs upstream of the translational start site of the gene encoding ELF4 (Li et al. 2011). Also, pairs of G-and N-boxes were found to be highly enriched in promoters targeted by the bHLH PIF1 (Kim et al. 2016). It is possible that multiple *cis*-elements serve to recruit additional TFs for *in vivo* cooperative binding.

### C_4_ photosynthesis co-opted an ancient C_3_ *cis*-regulatory code built on G-box recognition by bHLH transcription factors

Finally, from this work we propose a model that summarises how molecular evolution of *cis*-elements recognised by bHLHs may relate to the recruitment of *NADP-ME* into C_4_ photosynthesis. C_4_ photosynthesis is an excellent example of convergent evolution (Sage et al. 2011; Christin et al. 2013) as it has evolved independently over 60 times in angiosperms (Sage et al. 2011; Sage 2016) and at least 22 times in grasses (Grass Phylogeny Working Group II 2012). How this repeated evolution has come about is not fully understood. Our model contributes to our understanding of C_4_ evolution and is based on the following findings: first, in rice, which belongs to the BEP clade that contains no C_4_ species, only one copy of a G-box was present in the *NADP-ME* promoter. In contrast, *cis*-element pairs recognised by ZmbHLH128 and ZmbHLH129 in *NADP-ME* promoters seem to be common in the PACMAD clade that contains many independent C_4_ lineages. For example, in the PACMAD grasses a G-and N-box pair was identified in C_3_ *D. oligosanthes* (Do024386) and appears to be reasonably conserved in C_4_ species in this group. However, in the case of the C_4_-*NADP-ME*s from *S. bicolor* and *Z. mays* (Sobic.003g036200 and GRMZM2G085019) these elements have diversified. Both of these grass species belong to the C_4_ lineage Andropogoneae in which the plastidic NADP-ME isoform that is used in C_4_ photosynthesis (C_4_-NADP-ME) evolved by duplication from an ancestral plastidic NADP-ME that still exists and is not involved in the C_4_ cycle (*non*C_4_-NADP-ME, Sobic.009g108700 and GRMZM2G122479) (Tausta et al. 2002; Maier et al. 2011; Alvarez et al. 2013). In contrast, C_4_ *S. italica* together with C_3_ *D. oligosanthes* belong to the grass lineage Paniceae in which only one plastidic NADP-ME isoform is known to exist (Si000645 and Do024386) (Alvarez et al. 2013; Emms et al. 2016). Surprisingly, the *cis*-element pair identified in the C_4_-*NADP-ME* promoter from *S. italica* (G-and N-box) was found to be closer to those occurring in the C_3_ and *non*C_4_-*NADP-ME* promoters from *D. oligosanthes*, *S. bicolor*, and *Z. mays* (G-and N-box/N-box-like) than to those occurring in the C_4_-*NADP-ME* promoters from *S. bicolor* and *Z. mays* (FeRE1 and N-box or FBS and FBS, respectively). A similar trend has previously been observed (Alvarez et al. 2013) and may be explained by the independent evolutionary origin of C_4_ photosynthesis in grass lineages formed by *S. italica* (Paniceae) or *S. bicolor*/*Z. mays* (Andropogoneae).

Taken together, our findings suggest that an ancestral G-box in combination with N-box-derived *cis*-elements form the basis of the synergistic binding of either ZmbHLH128 or ZmbHLH129 to *NADP-ME* promoters from PACMAD grasses. Nucleotide diversity in *cis*-elements recognised by bHLH TFs has been suggested as one of the mechanisms by which these TFs are involved in complex and diverse transcriptional activity (Toledo-Ortiz et al. 2003). We, therefore, can not exclude the possibility that the gene encoding the plastidic NADP-ME from C_3_ BEP *Brachypodium distachyon* (BRADI2g05620) can also be bound by ZmbHLH128 or ZmbHLH129 despite none of the typical *cis-*elements recognised by bHLH being identified in the promoter. Given recent evidence indicating that the bHLH TF family is often recruited into C_4_ photosynthesis regulation (Huang and Brutnell 2016), we suggest that the observed nucleotide modifications in the *cis-*element pair present in C_4_-*NADP-ME* promoters from *S. bicolor* and *Z. mays* may underlie changes in bHLH binding specificity *in vivo* and, therefore, contribute to the *NADP-ME* recruitment into C_4_ photosynthesis in the Andropogoneae lineage from the PACMAD clade. The presence of a bHLH duplicate (ZmbHLH129) that seems not to be required for C_4_ photosynthesis and has evolved to repress the activity of its homeolog (ZmbHLH128) is unique to maize as this homeolog gene pair resulted from the maize WGD. Therefore, we hypothesise that the single orthologous bHLH in all the other PACMAD species activates C_4_-*NADP-ME* gene expression. This agrees with the hypothesis that C_4_ photosynthesis has on multiple occasions made use of *cis*-regulators found in C_3_ species and, therefore, that the recruitment of C_4_ genes was made through minor rewiring of pre-existing regulatory networks (Reyna-Llorens and Hibberd 2017). We conclude that regulation of C_4_ genes can be based on an ancient code founded on a G-box present in the BEP clade as well as the PACMADs. Acquisition of a second *cis*-element recognised by bHLH in the PACMAD clade appears to have facilitated synergistic binding by either ZmbHLH128 or ZmbHLH129. Although this G-box-based *cis*-code has remained similar in *S. italica*, it has diverged in maize and sorghum. Thus, different C_4_ grass lineages may employ slightly different molecular circuits to regulate orthologous C_4_ photosynthesis genes.

## Materials and methods

### Plant growth conditions and collection of leaf samples

To construct the cDNA expression library, maize plants (*Zea mays* L. var. B73) were grown at 16h photoperiod with a light intensity of 340-350 μmol m^−2^ s^−1^, at day/night temperature of 28°C/26°C, and 70% relative humidity. Two light regimes were used: (1) nine days in 16h photoperiod; and (2) nine days in 16h photoperiod followed by a 72h dark treatment. In both experiments, sample collection was performed under 16h photoperiod. Third leaves grown in the former and latter light regimes were harvested respectively at time points covering the Zeitgeber times (ZT) −0.5, 0.5, 2h, and ZT 1, 2, 4, 8, 12, 15.5h. For isolation of maize mesophyll protoplasts, maize plants were grown for 10 days at 25°C, 16h photoperiod (60 μmol m^−2^ s^−1^), and 70% relative humidity. For transient expression assays *in planta, Nicotiana benthamiana* (tobacco) plants were grown for five weeks at 22°C, 16h photoperiod (350 μmol m^−2^ s^−1^), and 65% relative humidity. After agro-infiltration of tobacco leaves, plants were left to grow into the same growth conditions and leaf discs (2.5 cm in diameter) collected 96h post-infection.

### Generation of yeast bait strains

Yeast bait strains were generated as previously described (Ouwerkerk and Meijer 2001; Serra et al. 2013). Yeast strain Y187 (Clontech) was used to generate six bait strains carrying overlapping fragments of the *ZmC_4_-NADP-ME* (GRMZM2G085019) promoter cloned into the yeast integrative vector pINT1-HIS3 (Ouwerkerk and Meijer 2001) as *Not*I-*Spe*I or *Xba*I-*Spe*I fragments (supplementary table S1). The *ZmC_4_-NADP-ME* promoter region was defined as the 1982 bp upstream of the predicted translational start site (ATG). To assess self-activation/*HIS3* leaky expression, yeast bait strains were titrated in complete minimal medium (CM) lacking histidine, with increasing concentrations of 3-amino-1,2,4-triazole (3-AT, up to 75 mM).

### Construction of cDNA expression library

Total RNA was extracted from third leaves of maize seedlings using TRIzol reagent (Invitrogen), following the manufacturer’s instructions. RNA samples from nine time points (described in ‘plant growth conditions and collection of leaf samples’) were pooled in equal amounts for mRNA purification using the PolyATract mRNA Isolation System IV (Promega). A unidirectional cDNA expression library was prepared using the HybriZAP-2.1 XR cDNA Synthesis Kit and the HybriZAP-2.1 XR Library Construction Kit (Stratagene), following the manufacturer’s instructions. Four micrograms of mRNA were used for first strand cDNA synthesis. After *in vivo* excision and amplification of the pAD-GAL4-2.1 phagemid vector, this maize cDNA expression library was used to transform yeast bait strains.

### Yeast One-Hybrid (Y1H) screening and validation

Yeast bait strains were transformed with 1 μg of maize cDNA expression library according to Ouwerkerk and Meijer (2001) and Serra et al. (2013). At least, 1.3 million yeast colonies of each yeast bait strain transformed with the maize cDNA expression library were screened in CM-HIS-LEU supplemented with 3-AT: 5 mM (-1982 to −1524 bp), 20 mM (-389 to −154 bp, −776 to −334 bp) or 75 mM (-973 to −702 bp, −1225 to −891 bp, −1617 to −1135 bp). Plasmids from yeast clones that actively grew on selective medium were extracted. To know whether the isolated clones encoded transcription factors (TFs), the cDNA insert was sequenced and the results analysed using BLAST programmes. To validate DNA-TF interactions in yeast, isolated plasmids encoding TFs were re-transformed into the yeast bait strain in which they were found to bind. To assess TF binding specificity, plasmids encoding TFs were also transformed into the yeast bait strains to which they do not bind.

### Yeast cell spotting

Yeast bait strains transformed with plasmids encoding TFs were grown overnight until log or mid-log phase at 30°C in liquid yeast CM medium supplemented with Histidine (CM +HIS-LEU). Cultures were normalized to an OD_600_ of 0.4, spotted onto solid medium CM +HIS-LEU or CM-HIS-LEU + 3-AT, and grown for 3 days at 30°C.

### Isolation and transformation of maize mesophyll protoplasts

Maize mesophyll protoplasts were isolated from 10-day-old maize greening plants and transformed according to Lourenço et al. (2013) with minor modifications. Mid-section of newly matured second leaves was digested in a cell wall digestive medium containing 1.5% (w/v) cellulase R-10 (Duchefa), 0.3% (w/v) macerozyme R-10 (Duchefa), 10 mM MES (pH 5.7), 0.4 M mannitol,1 mM CaCl_2_, 0.1% (w/v) BSA and 5 mM β-mercaptoethanol. Several leaf blades were stacked and cut perpendicularly to the long axis into 0.5 to 1 mm slices and quickly transferred to digestive medium (25 mL digestive medium for each set of 10 leaf blades). Purity and integrity of isolated protoplasts were examined under light microscopy. Mesophyll protoplasts were quantified and its abundance adjusted to 2 × 10^6^ protoplasts/mL. Transformed protoplasts were resuspended in 1.25 mL of incubation solution (0.6 M mannitol, 4 mM MES (pH 5.7) and 4 mM KCl) and incubated in 24-well plates for 18h at room temperature under dark.

### Bimolecular Fluorescence Complementation (BiFC) assay

To generate BiFC constructs, full-length coding sequences (CDS) of *ZmbHLH128* (GRMZM2G314882) and *ZmbHLH129* (GRMZM5G856837) were PCR-amplified using respectively the following pairs of *att*B-containing primers: 5’-GGGGACAAGTTTGTACAAAAAAGCAGGCTNNATGATGAACTGCGCCGGA-3’ / 5’-GGGGACCACTTTGTACAAGAAAGCTGGGTNCTAAGCATTAGGCGGCCAG-3’, and 5’- GGGGACAAGTTTGTACAAAAAAGCAGGCTNNATGATGGACTGCGCTGGA-3’ / 5’-GGGGACCACTTTGTACAAGAAAGCTGGGTNCTAAGCATTTGGGGGCCAG-3’ (underlined sequences indicate *att*B Gateway adaptors). *ZmbHLH128* and *ZmbHLH129* CDS were recombined into pDONR221 (Invitrogen) to obtain Entry clones through BP-Gateway reaction (Invitrogen), following the manufacturer’s instructions. CDS were then recombined into vectors YFP^N^43 and YFP^C^43 through LR-Gateway reaction (Invitrogen) to raise a translational fusion with N-and C-terminal domains of yellow fluorescent protein (YFP), respectively. Final BiFC constructs were denominated as YFP^N^∷ZmbHLH128, YFP^N^∷ZmbHLH129, YFP^C^∷ZmbHLH128, and YFP^C^∷ZmbHLH129. Maize mesophyll protoplasts were transformed with 6 μg of each of the BiFC constructs. Protoplasts transformed with YFP^N^∷Akin10 (*Arabidopsis* SNF1 Kinase Homolog 10), YFP^C^∷ Akin3 (*Arabidopsis* SNF1 Kinase Homolog 3) and YFP^N^43 and YFP^C^43 empty vectors were used as negative controls. Transformations were performed in triplicate. YFP fluorescence and chlorophyll autofluorescence signals were observed under a confocal microscope (Leica SP5).

### Transient expression assays *in planta*

For the transient expression assays in tobacco leaves, reporter and effector constructs were generated in the Gateway binary vectors pGWB3i (pGWB3 containing an intron-tagged β-glucuronidase (GUS) open reading frame (Berger et al. 2007)) and pGWB2 (Tanaka et al. 2012), respectively.

To construct the reporter plasmids, promoter fragments of *ZmC_4_-NADP-ME* (GRMZM2G085019, from −389 to −154 bp) and *ZmnonC_4_-NADP-ME* (GRMZM2G122479, from −368 to −143 bp) were fused to a 136 bp minimal *CaMV35S* promoter (*m35S*) in a 3-step PCR reaction: (1) promoter sequences were amplified with long chimeric primers to introduce overlapping ends (reverse primer of *pZmC_4_-NADP-ME* / *pZmnonC_4_-NADP-ME* was designed to be complementary to the forward primer of the *m35S*) (supplementary table S4); (2) promoter sequences amplified by PCR in (1) were mixed according to the fusion products of interest in a ratio of 1:1 (*ZmC_4_-NADP-ME (-389 to −154 bp)∷m35S* and *ZmnonC_4_-NADP-ME(-368 to −143 bp)∷m35S*) and 10 PCR cycles were run without primers (denaturation at 98°C for 10 s, 55°C for 30 s, and 72°C for 1 min); and (3) fusion products of interest were amplified with *att*B-containing primers (supplementary table S4). To obtain Entry clones, promoter fragments fused to *m35S* were cloned into pDONR221 (Invitrogen) through BP-Gateway reaction (Invitrogen), following the manufacturer’s instructions. Promoter sequences were then recombined into the binary vector pGWB3i through LR-Gateway reaction (Invitrogen) to obtain the final reporter constructs for *promoter∷GUS* analysis (*pZmC_4_-NADP-ME* and *pZmnonC_4_-NADP-ME*). For the effector constructs (TF driven by the *CaMV35S* promoter), *ZmbHLH128* and *ZmbHLH129* Entry clones previously generated (see BiFC assay) were directly recombined into the binary vector pGWB2 through LR-Gateway reaction (Invitrogen).

Reporter and effector constructs together with a construct harbouring the silencing suppressor P1b (Valli et al. 2006) were transformed into the *Agrobacterium tumefaciens* strain GV301. Overnight cultures of Agrobacterium harbouring reporter, effector and P1b constructs were sedimented (5000 *g* for 15 min, at 4°C) and resuspended in infiltration medium (10 mM MgCl_2_, 10 mM MES (pH 5.6), 200 μM acetosyringone) to an OD_600_ of 0.3, 1, and 0.5, respectively, and mixed in a ratio of 1:1:1. Mixed Agrobacterium cultures were incubated for 2h at 28°C and used to spot-infiltrate the abaxial side of 5-week-old tobacco leaves. As controls, tobacco leaves were agro-infiltrated with mixed cultures carrying the reporter construct alone or the empty vector pGWB3i and effector constructs. Infected leaves were analysed at 96h post-infiltration. Leaf discs of 2.5 cm in diameter were collected from the infiltrated spots and used for the quantification of GUS activity. GUS activity was quantified by measuring the rate of 4-methylumbelliferyl-?-D-glucuronide (MUG) conversion to 4-methylumbelliferone (MU) as described in Jefferson et al. (1987) and Williams et al. (2016). Briefly, soluble protein was extracted from agro-infiltrated tobacco leaf discs by freezing in liquid nitrogen and maceration, followed by addition of protein extraction buffer. Diluted protein extracts (1:2) were incubated with 1 mM MUG for 30, 60, 90 and 120 min at 37°C in a 96-well plate. GUS activity was terminated at the end of each time point by the addition of 200 mM Na_2_CO_3_ and MU fluorescence measured by exciting at 365 nm and measuring emission at 455 nm. The concentration of MU/unit fluorescence in each sample was interpolated using a concentration gradient of MU from 1.5 to 800 μM MU.

### Production of recombinant ZmbHLH128 and ZmbHLH129

*ZmbHLH128* and *ZmbHLH129* full-length CDS were PCR-amplified using, respectively, the following pairs of gene specific primers 5’-GAATTCATGATGAACTGCGCCGGA-3’ / 5’-CTCGAGCTAAGCATTAGGCGGCCAG-3’ and 5’-GAATTCATGATGGACTGCGCTGGA-3’ / 5’-CTCGAGCTAAGCATTTGGGGGCCAG-3’ (underlined sequences indicate adaptors with restriction enzyme sites). *ZmbHLH128* and *ZmbHLH129* were cloned as *EcoR*I-*Xho*I fragments into the expression vector pET32a (Novagen), generating N-terminal Trx-tagged fusions. pET32a-Trx∷*ZmbHLH128* and pET32a-Trx∷*ZmbHLH129* constructs were confirmed by sequencing and transformed into Rosetta (DE3)pLysS competent cells (Invitrogen) for protein expression. Cells transformed with pET32a-Trx∷*ZmbHLH128* and pET32a-Trx∷*ZmbHLH129* constructs were respectively grown in Terrific Broth (TB) and Luria-Bertani (LB) medium to an OD_600_ of 0.5. Protein expression was induced with 4 mM isopropyl-d-1-thiogalactopyranoside (IPTG) and allowed to occur for 3h (ZmbHLH128) or 5h (ZmbHLH129) at 30°C. Protein purification was performed as described in Cordeiro et al. (2016).

### Blue Native-Polyacrylamide gel electrophoresis (BN-PAGE) and western blotting

Molecular mass of oligomers co-existing in purified ZmbHLH128 and ZmbHLH129 recombinant proteins was determined by blue native polyacrylamide gel electrophoresis (BN-PAGE). Two micrograms of the recombinant proteins (Trx∷His∷ZmbHLH128 or Trx∷His∷ZmbHLH129) were resolved on a 3-12% Novex Bis-Tris NativePAGE mini gel (Life Technologies), following the manufacturer’s instructions. HMW Native Marker Kit (66 - 669 kDa, GE Healthcare) was used to estimate molecular mass. Resolved proteins were transferred to a polyvinylidene difluoride (PVDF) membrane (GE Healthcare). The membrane was destained with a 50% (v/v) methanol and 10% (v/v) acid acetic solution followed by pure methanol. For immunodetection of Trx∷His∷ZmbHLH128 and Trx∷His∷ZmbHLH129, the membrane was incubated with α-His antibody (GE Healthcare) followed by α-mouse horseradish peroxidase-conjugated antibody (abcam) for 1h each at room temperature.

### Electrophoretic Mobility Shift Assay (EMSA)

DNA probes were generated by annealing oligonucleotide pairs in a thermocycler followed by radiolabeling as described in Serra et al. (2013). DNA probe sequences and respective annealing temperatures are listed in supplementary table S3. EMSAs were performed using 400 ng of the recombinant proteins Trx∷ZmbHLH128 or Trx∷ZmbHLH129, and 50 fmol of radiolabeled probes. Competition assays were performed adding 200- to 400-fold molar excess of the unlabeled probe. Trx∷OsPIF14 (LOC_Os07g05010) and Trx protein, both purified by Cordeiro et al. (2016), were used as negative controls. Each protein was mixed with probes in a 10 μl reaction containing 10 mM HEPES (pH 7.9), 40 mM KCl, 1 mM EDTA (pH 8), 1 mM DTT, 50 ng herring sperm DNA, 15 μg BSA and 10% (v/v) glycerol. Binding reactions were incubated for 1h on ice and the bound complexes resolved on a native 5% polyacrylamide gel (37.5:1). Gel electrophoresis and detection of radioactive signal were performed as described in Serra et al. (2013).

### Synteny analysis

SynFind (Tang et al. 2015) was used to identify maize syntenic chromosomal regions for *ZmbHLH128* (GRMZM2G314882) and *ZmbHLH129* (GRMZM5G856837) genes against *Z. mays* B73 RefGen_v3 genome. A table containing maize syntelog gene pairs was retrieved using SynFind tool (supplementary table S2).

### Phylogenetic analyses

*ZmbHLH128* and *ZmbHLH129* were used as references to identify closely related *bHLH* genes of *Zea mays*, *Sorghum bicolor*, *Setaria viridis*, *Setaria italica*, *Oryza sativa*, and *Brachypodium distachyon*, through Phytozome database (Goodstein et al. 2012). Predicted CDS were aligned using MUSCLE. The resulting alignment was used to infer a maximum likelihood phylogenetic tree, using GTR+G+I nucleotide substitution model (1000 bootstrap pseudoreplicates) in MEGA 7 software (Kumar et al. 2016). Phylogenetic analysis of genes encoding C_3_ and C_4_ plastidic NADP-ME isoforms from *B. distachyon* (BRADI2g05620), *O. sativa* (LOC_Os01g09320), *D. oligosanthes* (Do024386), *S. italica* (Si000645), *S. bicolor* (Sobic.003g036200, Sobic.009G108700) and *Z. mays* (GRMZM2G085019, GRMZM2G122479) was performed using Geneious Pro 5.3.6 software (Kearse et al. 2012). Full-length genomic sequences were aligned using MUSCLE. Phylogenetic tree was inferred using the Neighbor Joining (1000 bootstrap pseudoreplicates) and rooted using the gene encoding C_3_ plastidic NADP-ME (At1g79750) from *Arabidopsis thaliana*, a dicot angiosperm.

## Acknowledgments

We thank Lisete Fernandes (Escola Superior de Tecnologia da Saúde de Lisboa, Portugal) for discussions and advice concerning EMSA experiments, Cecília Arraiano Lab (ITQB-NOVA, Oeiras, Portugal) for material used in EMSA experiments, Myriam Goudet and Samuel Brockington (Department of Plant Sciences, University of Cambridge, UK) for assistance with phylogenetic analyses. Fundação para a Ciência e Tecnologia (FCT) is acknowledged through research unit GREEN-it ‘Bioresources for Sustainability’ (UID/Multi/04551/2013). ARB (SFRH/BD/105739/2014), AG (SFRH/BD/89743/2012), AMC (SFRH/BD/74946/2010), PMB (SFRH/BPD/86742/2012), IAA (IF/00960/2013 – POPH-QREN), and NJMS (IF/01126/2012 – POPH-QREN) were funded by FCT, TSS and PG by European Union project *3to4* (Grant agreement no: 289582), and IR-L by BBSRC grant (BB/L014130).

